## Supplementary Information for "Structural basis of inhibition of human Na_V_1.8 by the tarantula venom peptide Protoxin-I"

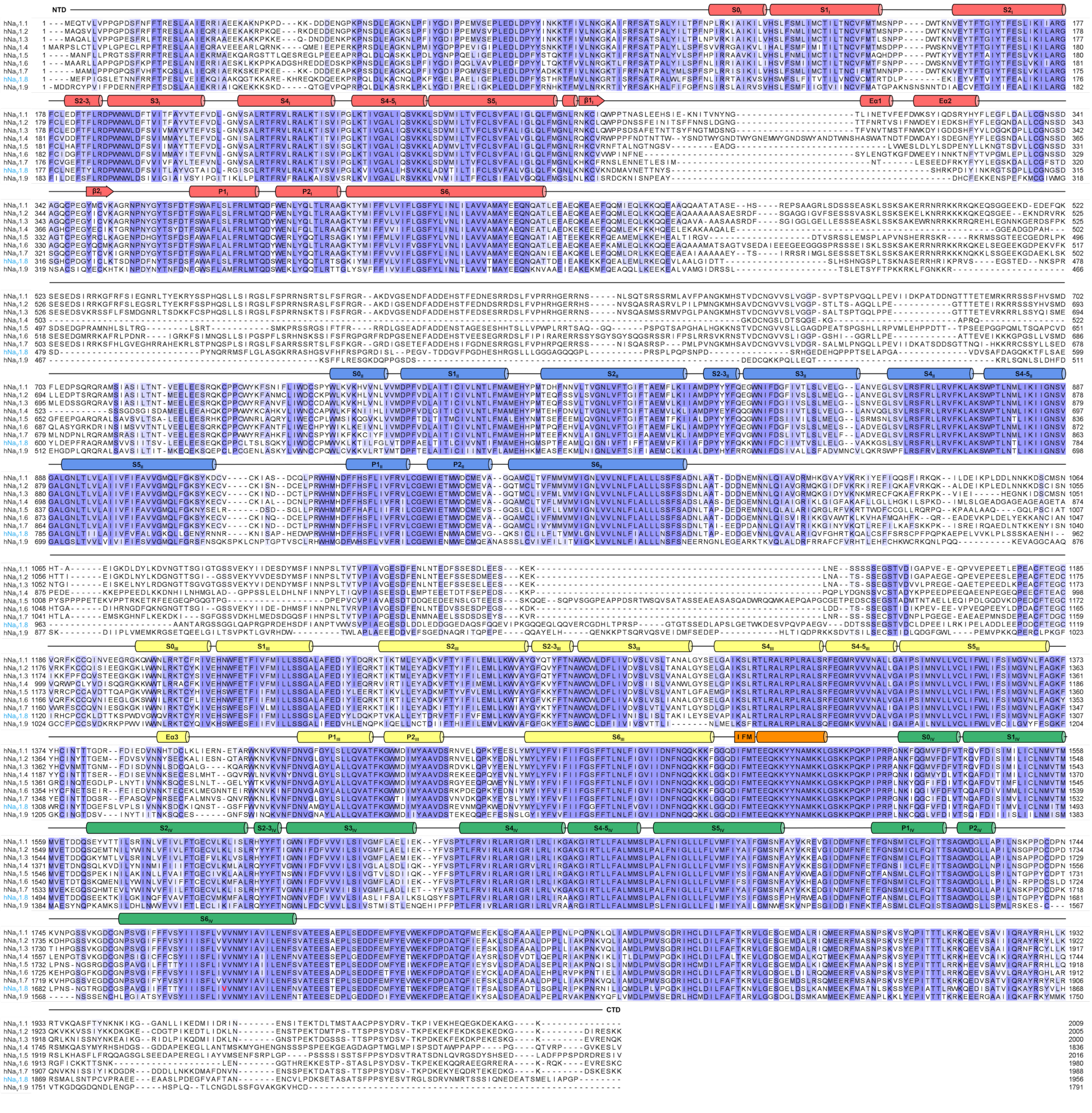

**Supplementary Figure 1: Sequence alignment of hNa$_V$1.1-1.9**

Sequences were aligned using ClustalΩ,[1] and viewed in Jalview.[2] Residues are colored by conservation score; domains are colored following the scheme in Figure **1a**

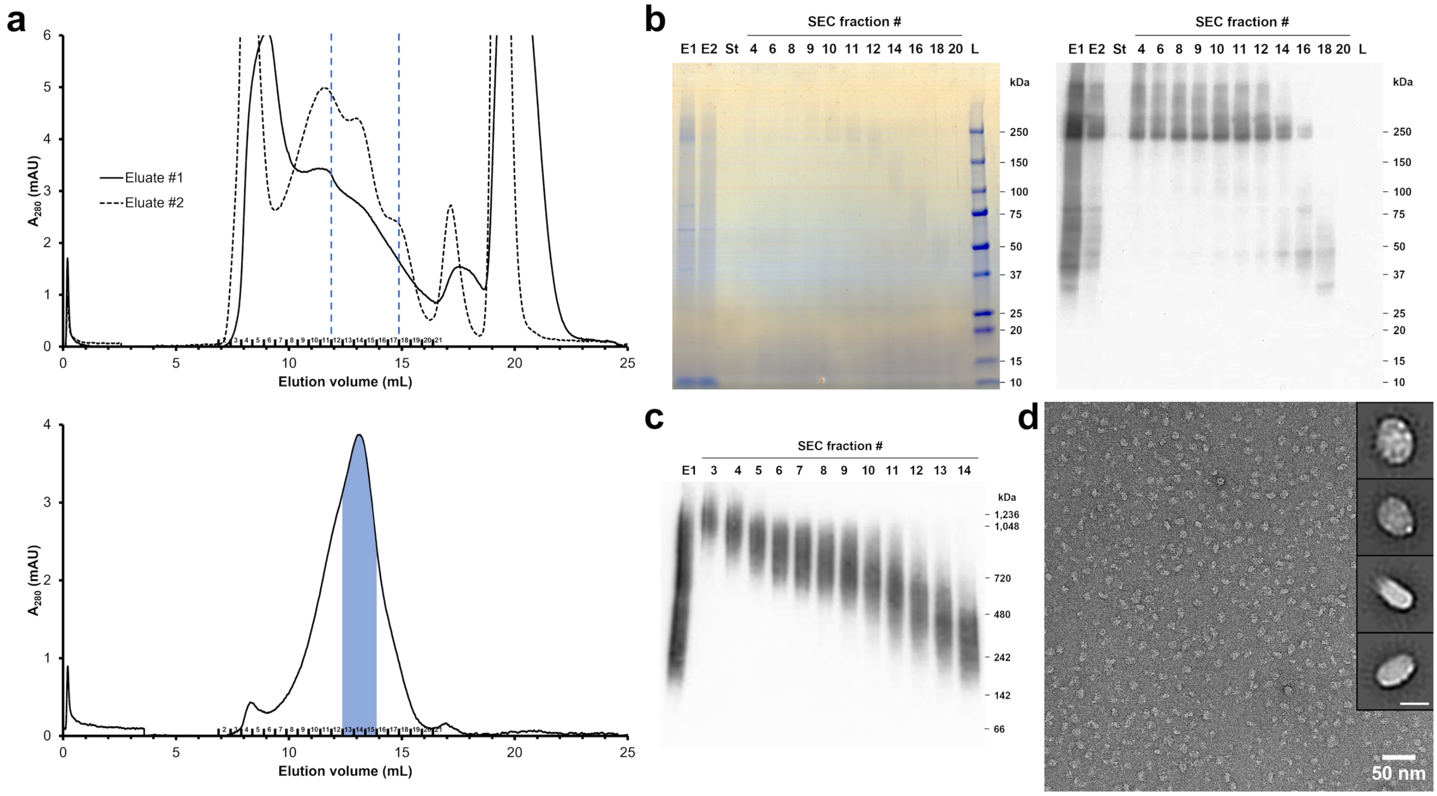


**Supplementary Figure 2: Biochemistry and purification of apo-hNa$_V$1.8**

**a** (upper) Example size-exclusion chromatography (SEC) traces following FLAG resin purification; the Eluate #1 trace shown as a solid black line and Eluate #2 as a dashed black line. Fractions between the dashed blue lines were pooled for (lower) subsequent SEC purification. Fractions highlighted in solid blue (F13-15) were carried forward for cryoEM. **b** SDS-PAGE gels of the fractions from the final SEC purification, with proteins detected by (left) Coomassie blue stain and (right) anti-FLAG western blotting. **c** Native PAGE gel of the fractions from the final SEC purification, with proteins detected by anti-FLAG western blotting. **d** Representative micrograph of negatively-stained particles from pooled fractions F13-15 with (inset) selected 2D class averages. Inset scale bar = 15 nm

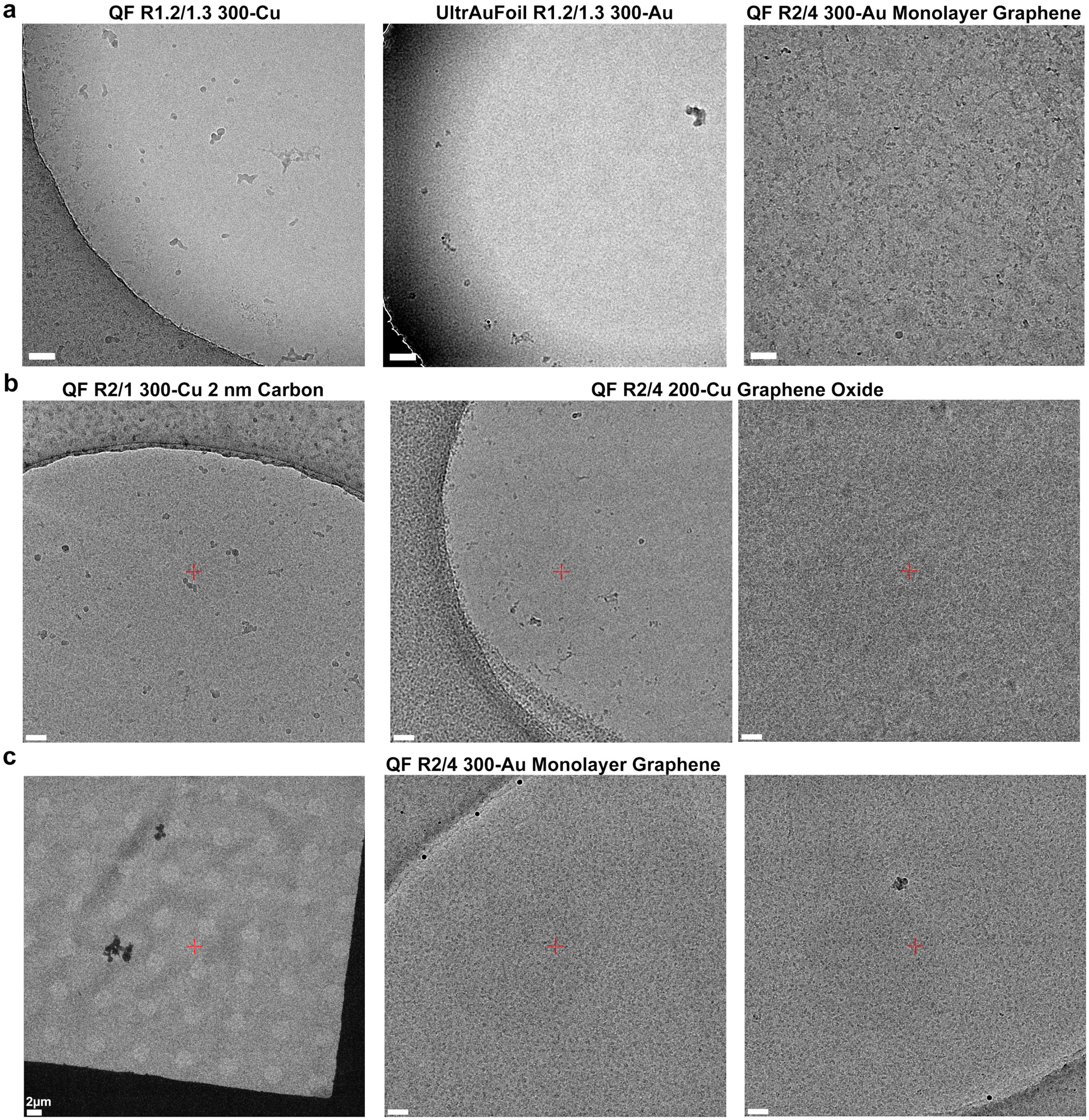


**Supplementary Figure 3: Grid freezing overview outlining particle distribution on grids with and without support films**

Representative cryoEM grid images of apo-hNa$_V$1.8. Scale bars are 50 nm unless otherwise indicated. **a** Example images of different grid types during initial screening; (left) Quantifoil R1.2/1.3 300-Cu grids show particles on carbon support rather than in holes; (center) UltrAuFoil R1.2/1.3 300-Au grids show particles in thick ice at the edge of the holes; and (right) Quantifoil R2/4 300-Au monolayer graphene grids show improved particle distribution. **b** Example images of different support film grids; (left) 2 nm carbon support film shows particles predominantly distributed on the ultrathin carbon support; (center) graphene oxide grids show frequent breakage after glow discharging even as (right) they show good particle distribution on the support film. **c** Example images of the grid leading to the apo-hNa$_V$1.8 reconstruction showing good particle distribution and contrast on the monolayer graphene support

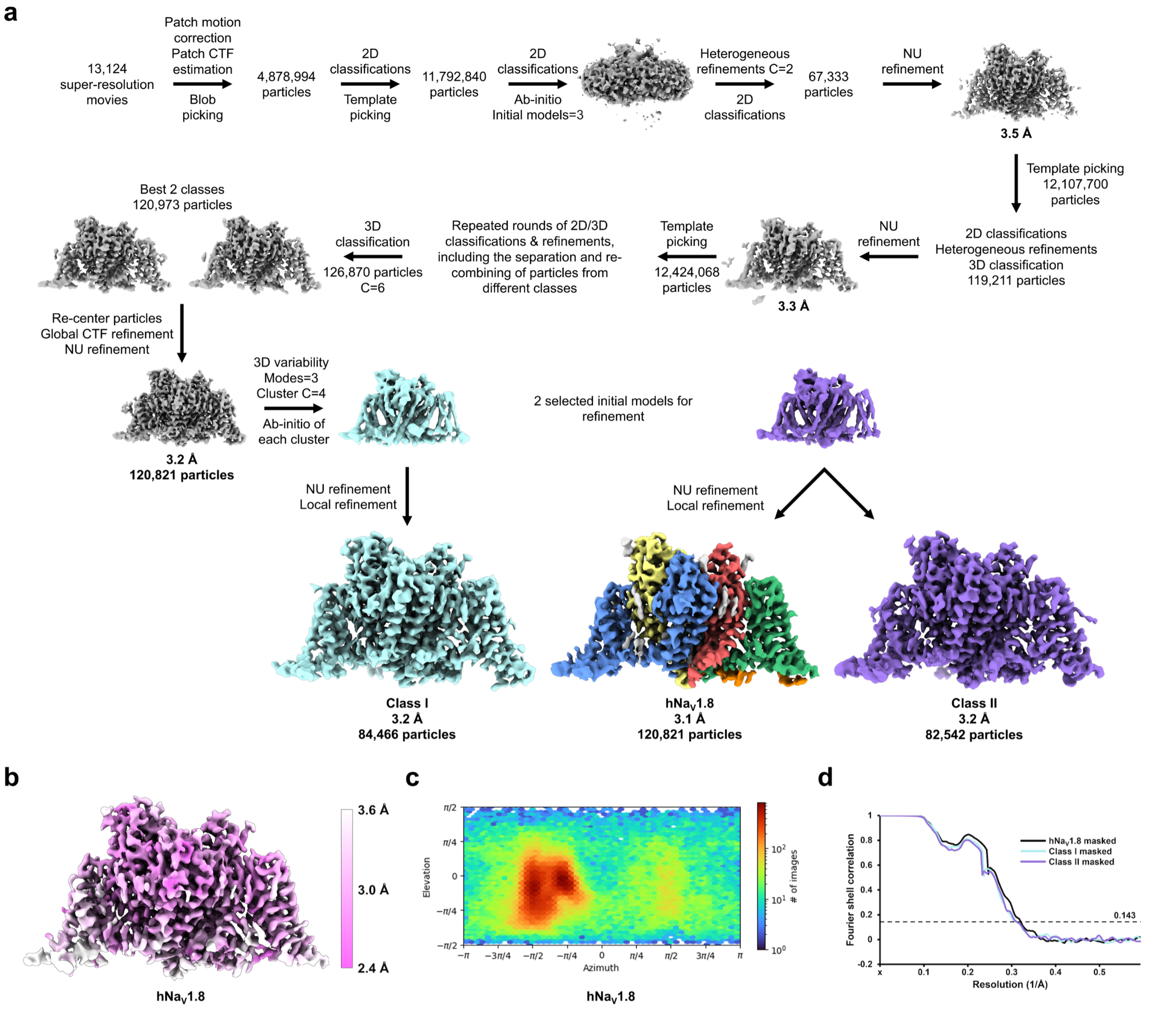


**Supplementary Figure 4: Processing flowchart for apo-hNa$_V$1.8 using CryoSPARC**

**a** Processing workflow resulting in the apo-hNa$_V$1.8 reconstruction, along with Class I and Class II. **b** Local resolution of apo-hNa$_V$1.8 calculated within CryoSPARC. **c** Angular distribution plot of particles in the final apo-hNa$_V$1.8 reconstruction. **d** Fourier shell correlation (FSC) plot of the three final apo-hNa$_V$1.8 reconstructions. FSC = 0.143 is indicated by a dashed black line

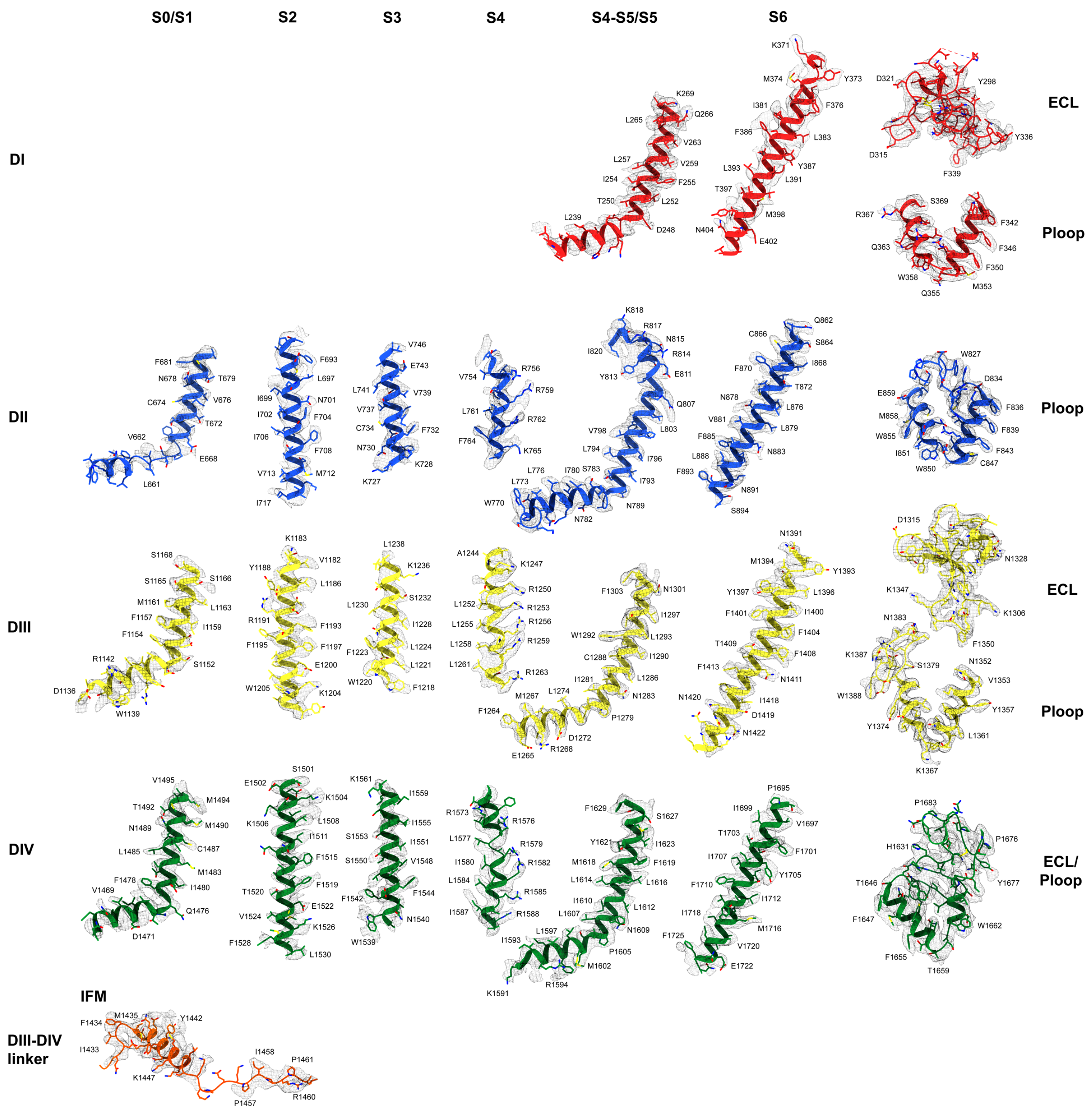


**Supplementary Figure 5: apo-hNa$_V$1.8 model-to-map fit**
All domains are colored according to the scheme in Figure **1a**

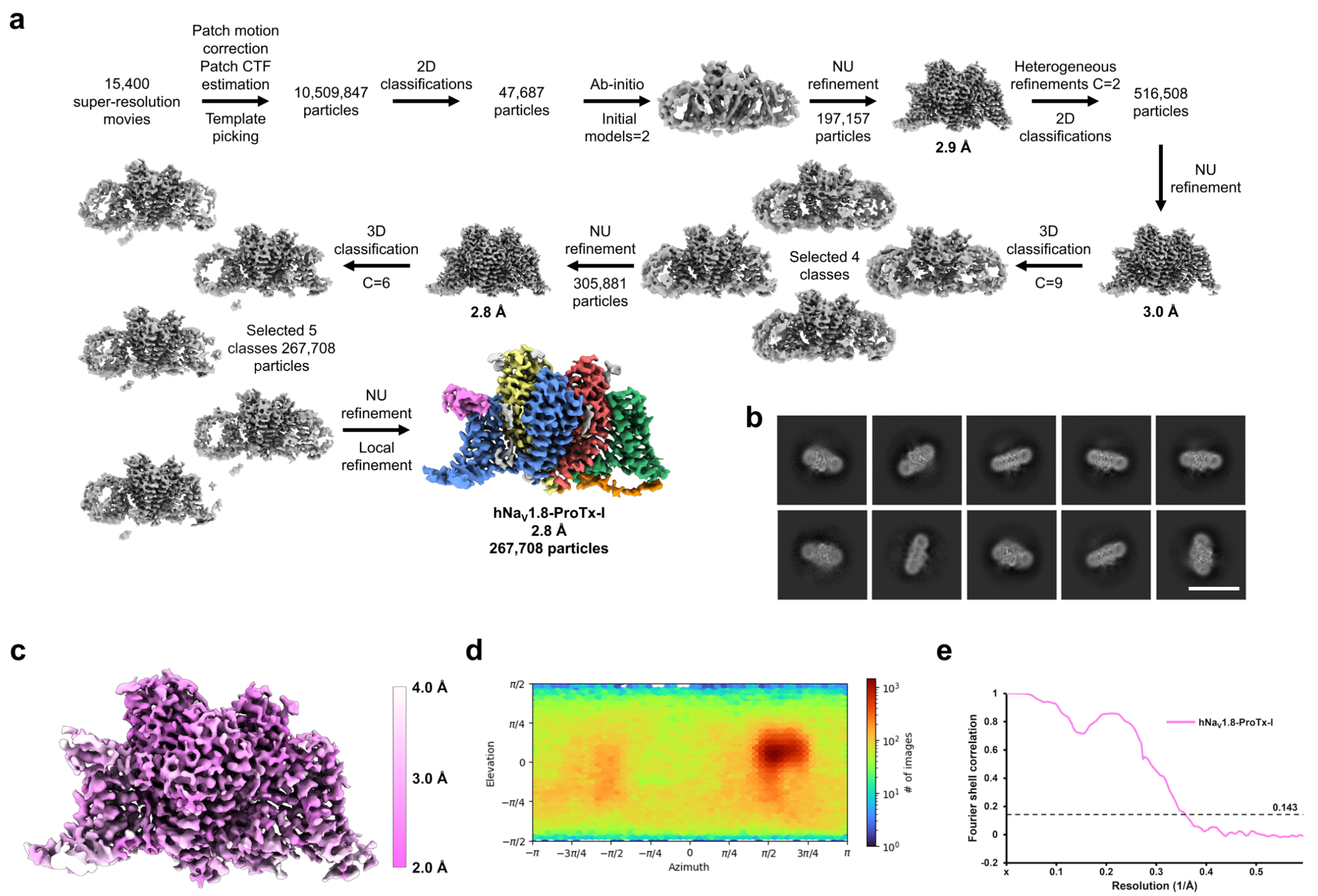


**Supplementary Figure 6: Processing flowchart for the hNa$_V$1.8-ProTx-I complex using CryoSPARC**
**a** Processing workflow resulting in the hNa$_V$1.8-ProTx-I reconstruction. **b** Example 2D class averages. Scale bar = 15 nm. **c** Local resolution of hNa$_V$1.8-ProTx-I calculated within CryoSPARC. **d** Angular distribution plot of particles in the final hNa$_V$1.8-ProTx-I reconstruction. **e** FSC plot of the final hNa$_V$1.8-ProTx-I reconstruction. FSC = 0.143 is indicated by a dashed black line

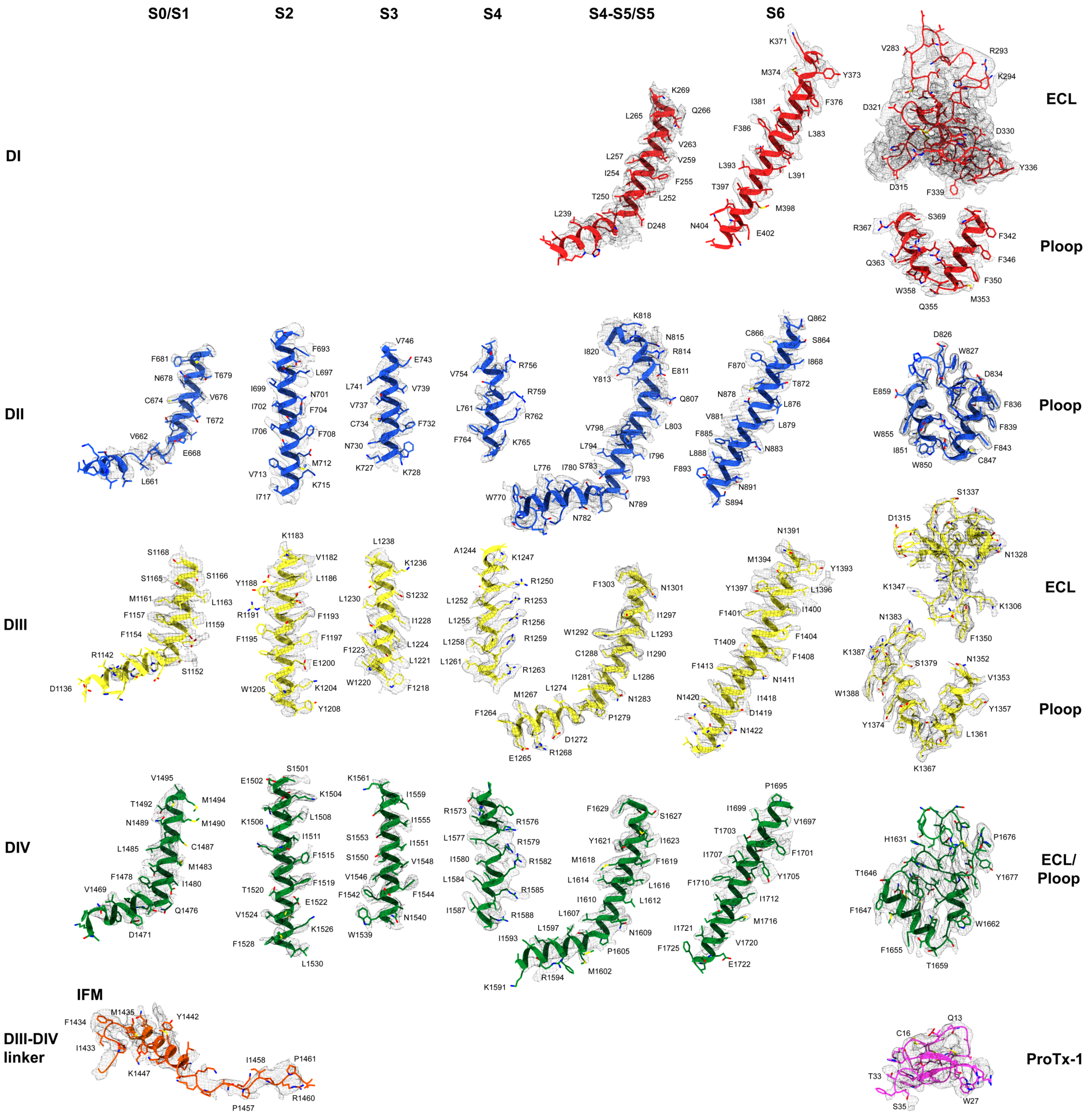


**Supplementary Figure 7: hNa$_V$1.8-ProTx-I model-to-map fit**
All domains are colored according to the scheme in Figure 1a, with ProTx-I colored in pink

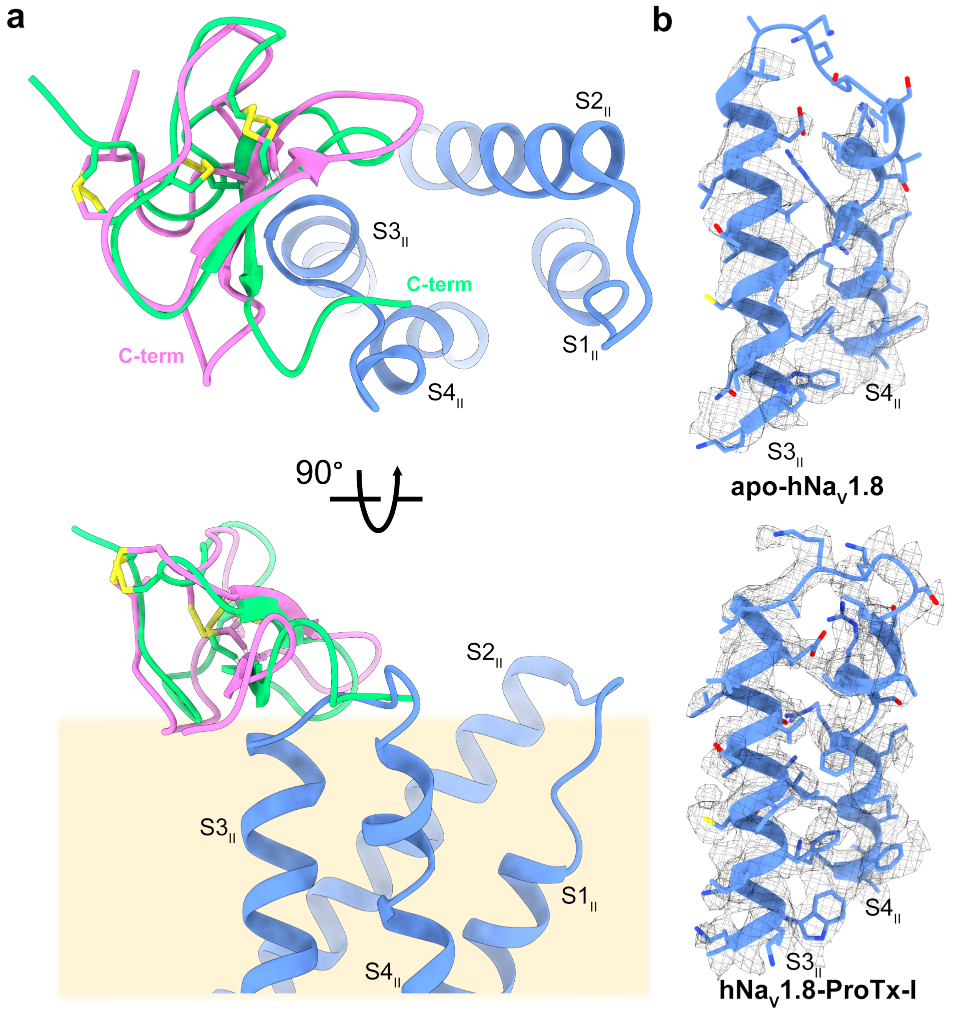


**Supplementary Figure 8: ProTx-I interactions and alignment with hNa$_V$1.8**

**a** Extracellular view of ProTx-I binding region highlighting the repositioning of the C-terminus. Bound ProTx-I from this study in pink and an NMR model of ProTx-I in green (PDB 2M9L). **b** Model-to-map fit of VSD$_{II}$ S3 and S4 helices for apo-hNa$_V$1.8 (top) and hNa$_V$1.8-ProTx-I (bottom) highlighting the observed comparative lower-resolution of the S3-S4 linker in apo-hNa$_V$1.8

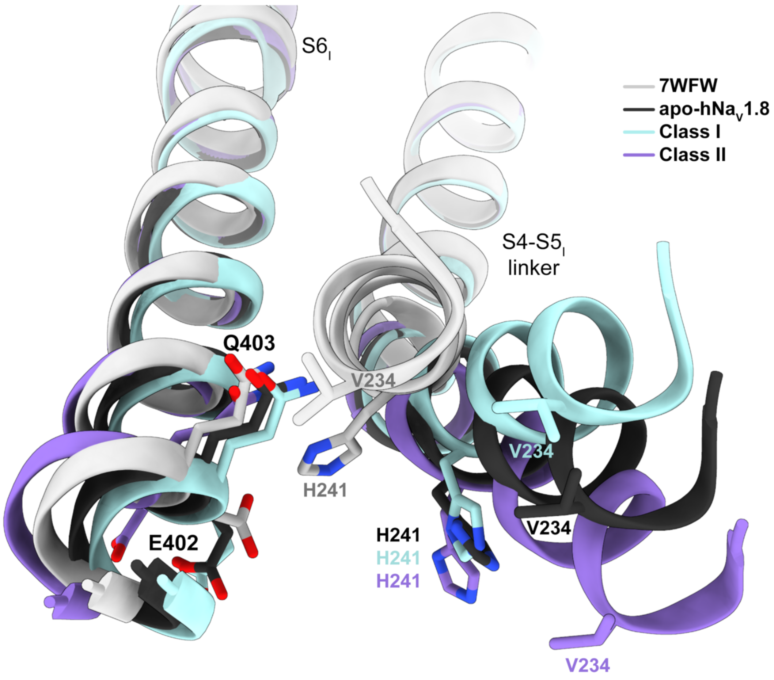


**Supplementary Figure 9: DI S4-S5 linker and S6 region depicting location of key residues**
Residues H241 and V234, unique to $hNa_V1.8$ are indicated, along with conserved residues E402 and Q403 on DI S6 helix. Domains are colored following the scheme in Figure 2a

**Supplementary Movie 1: 3D variability analysis prior to final reconstruction of apo-hNa$_V$1.8 highlighting the movements of VSD$_I$ S4-S5 linker**
Domains are colored following the scheme in Figure **1a**

**Supplementary Movie 2: 3D variability analysis prior to final reconstruction of apo-hNa$_V$1.8 highlighting the NTD**
Domains are colored following the scheme in Figure **1a**. NTD density on the bottom left in white.

**Supplementary Movie 3: Hydrophobic surface of ProTx-I from the hNa$_V$1.8-ProTx-I complex**
Colors follow the scheme in Figure **4a**

**Supplementary Movie 4: apo-hNa$_V$1.8 to hNa$_V$1.8-ProTx-I model morph highlighting the movement of the VSD$_{II}$ S3-S4 linker due to ProTx-I binding**
Apo-hNa$_V$1.8 is depicted in black; hNa$_V$1.8-ProTx-I in blue with ProTx-I in pink

| | apo-hNa$_V$1.8 | Class I | Class II | hNa$_V$1.8-ProTx-I |
|---|---|---|---|---|
| **Data collection** | | | | |
| Microscope | Titan Krios | | | |
| Voltage (kV) | 300 | | | |
| Detector | Gatan K3 | | | |
| Pixel size (Å) | 0.839 | | | 0.827 |
| Total electron dose ($e^-$/Å$^2$) | 60 | | | |
| Defocus range (µm) | –1 to –2.5 | | | –1 to –2 |
| # of movies | 13,124 | | | 15,400 |
| **Reconstruction** | | | | |
| Software | CryoSPARC | | | |
| Symmetry | C1 (no symmetry) | | | |
| Selected movies | 13,027 | | | 14,125 |
| Final # of particles | 120,821 | 84,466 | 82,542 | 267,708 |
| Overall resolution (Å) | 3.12 | 3.24 | 3.22 | 2.76 |
| FSC threshold | 0.143 | | | |
| Map sharpening B factor (Å$^2$) | -73.97 | -68.94 | -64.86 | -65.68 |
| **Model Refinement** | | | | |
| Model composition | | | | |
| Non-hydrogen atoms | 8,234 | 8,234 | 8,245 | 8,638 |
| Protein residues | 998 | 998 | 998 | 1047 |
| Ligands | 14 | 14 | 15 | 15 |
| R.M.S. deviations | | | | |
| Bond lengths (Å) | 0.005 | 0.006 | 0.006 | 0.006 |
| Bond angles (°) | 0.758 | 0.831 | 0.874 | 0.957 |
| Validation | | | | |
| MolProbity score | 1.67 | 1.82 | 1.75 | 1.97 |
| Clashscore | 7.82 | 9.08 | 8.89 | 9.37 |
| Poor rotamers (%) | 0.45 | 0.79 | 0.68 | 1.72 |
| Ramachandran plot (%) | | | | |
| Favored | 96.36 | 95.15 | 96.06 | 95.77 |
| Allowed | 3.64 | 4.85 | 3.94 | 4.23 |
| Outlier | 0.00 | 0.00 | 0.00 | 0.00 |

**Supplementary Table 1: Data collection parameters, model statistics and validation**
